# Absence of a spindle position checkpoint in the fungal pathogen *Cryptococcus neoformans*

**DOI:** 10.64898/2026.08.31.748335

**Authors:** Taylor K. Wang, Julia Matthews, Abigail Strege, Edward W.J. Wallace, Xiaoxue Zhou

## Abstract

To maintain genome stability, it is crucial that cells do not initiate cytokinesis until chromosomes have been properly segregated. In the model budding yeast *Saccharomyces cerevisiae*, a surveillance mechanism called the Spindle Position Checkpoint (SPoC) ensures this coordination by regulating the Mitotic Exit Network (MEN) to couple exit from mitosis and cytokinesis to spindle position. The MEN is conserved in Ascomycota where the orthologous pathway in the fission yeast *Schizosaccharomyces pombe*, the Septation Initiation Network (SIN), regulates cytokinesis in response to defects in spindle elongation. Here, we show that the MEN/SIN pathway is conserved in the basidiomycetous budding yeast and human pathogen, *Cryptococcus neoformans*, and controls cytokinesis. However, spindle position or elongation does not regulate pathway activation or cell cycle progression in *C. neoformans*. In essence, there appears to be no SPoC in this organism to delay cytokinesis upon defects in mitosis. We speculate that while increasing the risk of genome instability, the lack of a SPoC might facilitate *C. neoformans*’s ability to change ploidy in the host.

## INTRODUCTION

Fungal infections are a pressing and underappreciated threat to global public health, causing an estimated 2.5 million deaths each year^1^. One major contributing agent is *Cryptococcus neoformans*, a globally distributed opportunistic fungal pathogen ranked by the World Health Organization as “critical priority”^2^. During infection, *C. neoformans* undergoes fascinating morphological transitions including ploidy changes that play a major role in its pathogenicity and drug resistance^3–6^. Cell cycle alterations have been implied to underlie these morphological changes^7,8^. As a basidiomycetous budding yeast which diverged from the model budding yeast *Saccharomyces cerevisiae*, an ascomycete, more than 600 million years ago^9^ and evolved budding independently^10^, *C. neoformans* cell cycle control has both conserved and distinct features compared to *S. cerevisiae*^11–16^ with many aspects uncharacterized even under normal growth conditions. To understand its strategies in safeguarding genome integrity during mitotic divisions, we investigated a critical cell cycle checkpoint for budding yeasts, the spindle position checkpoint, in *C. neoformans*.

Cell division (cytokinesis) and nuclear division (mitosis) must be coordinated both temporally and spatially in eukaryotic organisms to maintain genome stability. Distinct strategies to achieve this coordination have been described across organisms^17,18^. In animal cells, the site of cytokinesis (division plane) is set by the mitotic spindle in late anaphase thus ensuring spatiotemporal coordination between mitosis and cytokinesis. In contrast, fungi and plants often specify the cell division site before the formation of the spindle. As a result, the mitotic spindle must be positioned appropriately to ensure nuclear division occurs across the predetermined division plane. This is best understood in the model budding yeast *S. cerevisiae* where the division plane (bud site) is set in G1, and two partially redundant pathways help position the spindle: a Kar9-dependent pathway that orients the spindle along the mother-bud axis and near the bud neck before anaphase^19,20^ and a dynein-based pathway that pulls one end of the spindle into the bud during anaphase^21,22^.

To ensure correct spindle positioning, a surveillance mechanism called the Spindle Position Checkpoint (SPoC) regulates the Mitotic Exit Network (MEN) to couple cell cycle progression to spindle position in *S. cerevisiae*^23–25^. The MEN is a GTPase-kinase signaling cascade that controls the M to G1 transition known as exit from mitosis (when the mitotic spindle is disassembled and chromosomes decondense) as well as cytokinesis in response to spindle position^26,27^. Only when one end of the spindle is delivered to the bud in anaphase indicating correct spindle position would the MEN be activated to trigger mitotic exit and cytokinesis^28^ (Fig. 1A). If the anaphase spindle remains entirely in the mother cell, the MEN is kept inactive which arrests the cell in anaphase and blocks cytokinesis until spindle position is corrected to prevent the production of anucleate buds.

**Figure 1.**
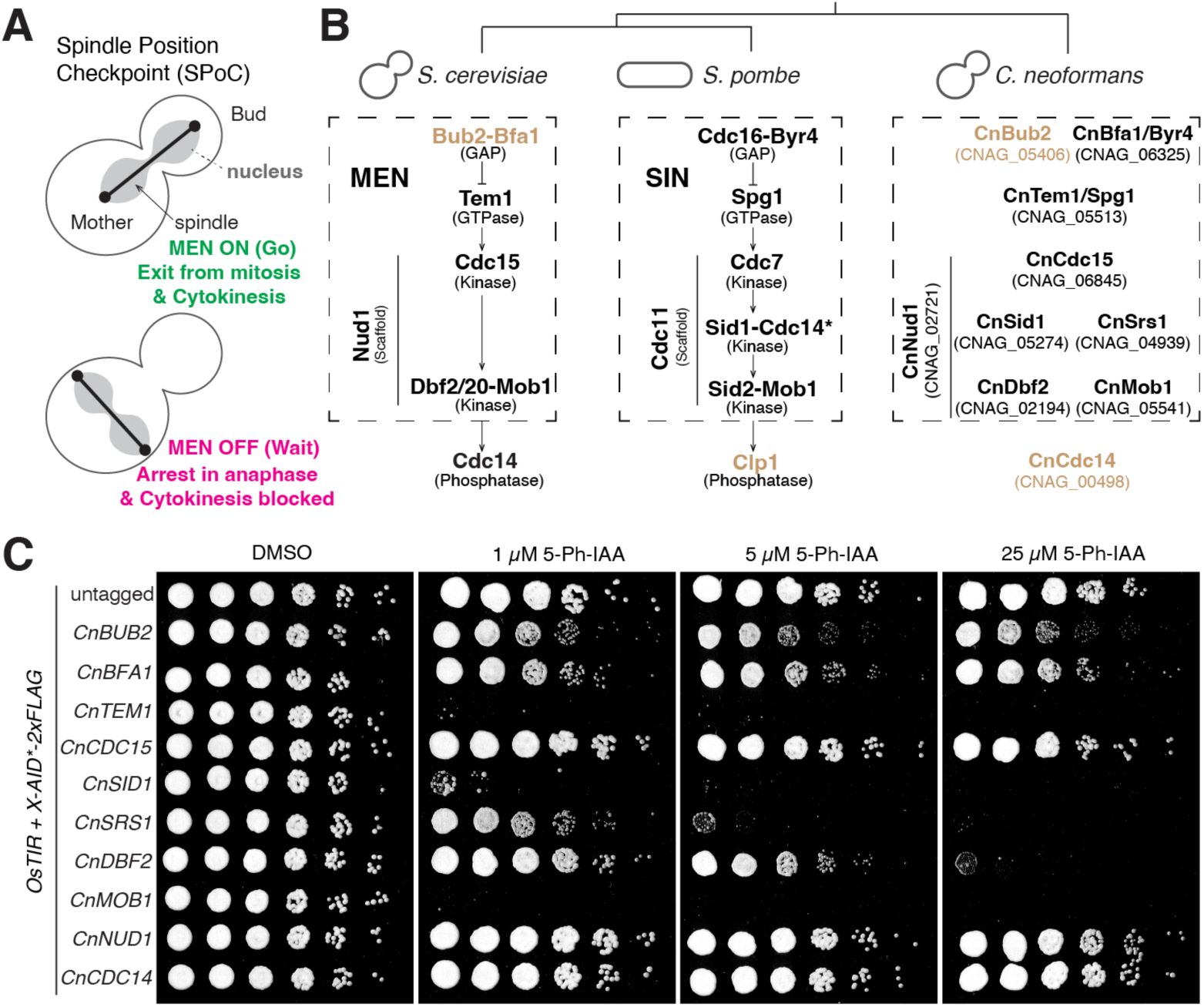
The core MEN/SIN pathway genes are present and essential in *C. neoformans*. (A) Illustration of the spindle position checkpoint (SPoC) in the model budding yeast *S. cerevisiae*. To produce viable progeny and ensure genome stability, only when the spindle is properly positioned across the bud neck in anaphase would the MEN pathway be activated to trigger exit from mitosis and cytokinesis (Go! signal). If the spindle is mispositioned, the MEN is kept inactive which arrests the cells in anaphase with cytokinesis blocked (Wait! signal) until the spindle is properly positioned. (B) Components of the MEN and SIN pathway and the corresponding orthologs we identified in *C. neoformans*. Black color denotes essential genes while the yellow color denotes nonessential genes. CnBfa1/Byr4 was determined to be essential in one study and predicted to be nonessential in another study. *, Cdc14 in *S. pombe* is a different gene from Cdc14 in *S. cerevisiae*. Genetic interaction arrows were omitted for *C. neoformans* as pathway topology has not been verified. (C) Growth analysis for auxin-inducible degradation of the conserved MEN/SIN core pathway components in *C. neoformans*. Serial dilutions of Cn26 (untagged), Cn53 (*CnBUB2-AID*), Cn45 (*CnBFA1-AID*), Cn35 (*CnTEM1-AID*), Cn37 (*CnCDC15-AID*), Cn39 (*CnSID1-AID*), Cn72 (*CnSRS1-AID*), Cn43 (*CnDBF2-AID*), Cn47 (*CnMOB1-AID*), Cn41 (*CnNUD1-AID*), and Cn49 (*CnCDC14-AID*) were spotted onto YPD with DMSO or increasing concentrations of 5-Ph-IAA and grown for 1.5 days at 30 °C.

The MEN has ancient roots in eukaryotes as orthologs of pathway components have been found in many eukaryotes. Studies in other ascomycetes ranging from budding^29,30^ and fission^31^ yeasts to filamentous fungi^32–34^ suggest that this pathway has a conserved function in regulating cytokinesis/septation in this major fungal phylum regardless of lifestyle, and a minor role in regulating mitotic exit in some organisms. In the limited cases where upstream signal of the pathway has been characterized, its role in coordinating cytokinesis with nuclear division as part of a spindle positioning checkpoint appears to be conserved. In the fungal pathogen *Candida albicans*, an ascomycete fungus that can grow as hyphae or yeast cells, a similar spindle position checkpoint delays cell cycle progression and cytokinesis upon spindle mispositioning during the yeast growth phase^35^. Like in *S. cerevisiae*, hyperactivation of the pathway by mutating the negative regulator (*bub2Δ*) bypasses the checkpoint and results in anucleate buds and multinucleate mothers when nuclear migration is inhibited. In the fission yeast *Schizosaccharomyces pombe*, the orthologous pathway known as Septation Initiation Network (SIN) regulates cytokinesis/septation in response to spindle elongation to ensure the delivery of nuclei to the cell poles before cytokinesis^36^.

Very little is known about the MEN/SIN pathway and the spindle position checkpoint in non-ascomycete fungi such as the basidiomycetes, which *C. neoformans* belongs to. Despite both dividing via budding, mitosis differs considerably in *C. neoformans* compared to *S. cerevisiae*. While *S. cerevisiae* undergoes closed mitosis with chromosomes segregating from the mother cell to the bud, *C. neoformans* undergoes semi-open mitosis with the replicated chromosomes first migrating to the bud and then half segregating back to the mother cell^11^. Intriguingly, in the corn smut *Ustilago maydis*, a dimorphic basidiomycetous fungus, orthologs of the MEN/SIN pathway were found to regulate nuclear envelope breakdown during mitosis^37^ suggesting a potential functional divergence of the pathway between Ascomycota and Basidiomycota. Thus, how basidiomycetous budding yeasts coordinate cytokinesis and mitosis within the spatiotemporal constraint of budding remains elusive.

Here we show that the MEN/SIN pathway is conserved in *C. neoformans* and controls cytokinesis just as in ascomycetes fungi. However, spindle position does not regulate the pathway activation or cell cycle progression. In essence, there does not appear to be a spindle position checkpoint in this organism. As a result, *C. neoformans* cells endure a higher risk of genome/ploidy instability upon exposure to microtubule poisons. We speculate that the lack of a spindle position checkpoint facilitates *C. neoformans*’s ability to undergo ploidy changes in the host which has been linked to its success as a pathogen.

## RESULTS

### The core MEN/SIN pathway genes are present and essential in C. neoformans

Since the MEN/SIN pathway has never been characterized in *C. neoformans*, we first checked whether the pathway is conserved by looking for pathway gene orthologs. While the core GTPase-kinase pathway topology is almost identical between the MEN and SIN, the SIN has an additional intermediate kinase complex (Fig. 1B), Sid1-Cdc14 (note that Cdc14 in *S. pombe* is not orthologous to Cdc14 in *S. cerevisiae*, which is a downstream target of the MEN). Recent studies suggest that the SIN composition is more ancestral and Sid1 was lost in *S. cerevisiae*^29^. We identified all the corresponding orthologs of core MEN/SIN genes in *C. neoformans* (Fig. 1B) including CnSid1 and its partner, the ortholog of *S. pombe* Cdc14, which we named CnSrs1 (for <u>S</u>id1 regulatory <u>s</u>ubunit) following the *C. albicans* gene name^30^ to avoid confusions with CnCdc14, the ortholog of *S. cerevisiae* Cdc14.

Consistent with the essential role of core MEN/SIN genes in *S. cerevisiae* and *S. pombe*, their orthologs in *C. neoformans* have also been classified as essential genes by previous studies^38–40^. However, there are two notable differences among the three organisms, namely the negative regulator of the pathway, the GAP complex, and the downstream effector phosphatase Cdc14/Clp1 (Fig. 1B). The GAP complex Bub2-Bfa1 is nonessential in *S. cerevisiae* albeit GAP mutants are SPoC defective, while the orthologous Cdc16-Byr4 is essential in *S. pombe* for preventing premature pathway activation. CnBub2 was designated as nonessential while its adaptor/scaffold CnBfa1/Byr4 was reported as nonessential by one study^39^ and essential by another^40^.

To validate the gene essentiality and to characterize the essential genes in the pathway, we adopted the auxin-inducible degron system (AID2) recently developed in *C. neoformans*^41^. After successfully fusing the degron (AID*-2xFLAG) at the C-terminus for each of the core pathway genes at the endogenous locus, we first confirmed the degradation upon auxin/ligand (5-Ph-IAA) addition (Fig. S1A). Next, we examined the effect of gene depletion on cell growth by plating cells on YPD agar with increasing concentrations of 5-Ph-IAA (Fig. 1C). Interestingly, genes in the pathway exhibited differential dosage sensitivity for 5-Ph-IAA (e.g., compare kinase CnSid1 with its partner CnSrs1, and the kinase CnDbf2 with its cofactor CnMob1). Nevertheless, we were able to confirm the essentiality for CnTem1, CnSid1, CnSrs1, CnDbf2 and CnMob1. We did not observe any growth defect for cells with the kinase CnCdc15 or the scaffold CnNud1 depleted at any of the ligand concentrations we tested despite both being designated as essential genes by previous studies. We speculate that the lack of growth defect upon protein depletion we observed is likely due to the AID2 system not being able to completely eliminate the expression of tagged proteins.

For the GAP complex CnBub2 and CnBfa1, we observed a considerable growth defect upon gene depletion while no growth defect was observed for CnCdc14 depletion (Fig. 1C). To confirm their non-essentiality, we constructed gene knockouts. We successfully generated knockouts strains for *CnBUB2* and *CnCDC14* which showed similar growth profiles as the AID knockdowns (Fig. S1C). However, multiple attempts to knockout *CnBFA1* were unsuccessful so we could not exclusively determine its gene essentiality.

We concluded that the core MEN/SIN pathway genes are present in *C. neoformans* and their gene essentiality is mostly conserved with some species differences where the GAP component CnBub2 is nonessential as in *S. cerevisiae* and the phosphatase CnCdc14 is nonessential as in *S. pombe*.

### The MEN/SIN pathway signals in a similar manner in *C. neoformans* via a GTPase signaling cascade that assembles on the spindle pole bodies

The MEN/SIN pathway is a GTPase-kinase signaling cascade^42^ that assembles on the spindle pole bodies (SPBs), which are microtubule-organizing centers (MTOCs) in fungi, for signal transduction^31,43,44^. To determine whether the identified pathway components in *C. neoformans* function in a similar manner, we first examined how the nucleotide state of the GTPase CnTem1 influences its function. To this end, we generated the corresponding GTP-locked (CnTem1-Q87L) and GDP-locked (CnTem1-T42A) mutants of CnTem1 and tested their abilities to support cell growth when the endogenous protein was depleted via the degron system. Like Tem1 in *S. cerevisiae*^42,45^ and Spg1 in *S. pombe*^46^, the GDP-locked mutant (T42A) did not complement CnTem1 depletion confirming the requirement for GTP-binding in signal transduction (Fig. 2A). Similar to the MEN in *S. cerevisiae* but unlike the SIN in *S. pombe*, hyperactivating the GTPase with the GTP-locked mutant (Q87L) was not lethal, but displayed a growth defect especially when wildtype Tem1 was depleted. This is consistent with our finding that CnBub2, the GAP for the GTPase, is not essential.

**Figure 2.**
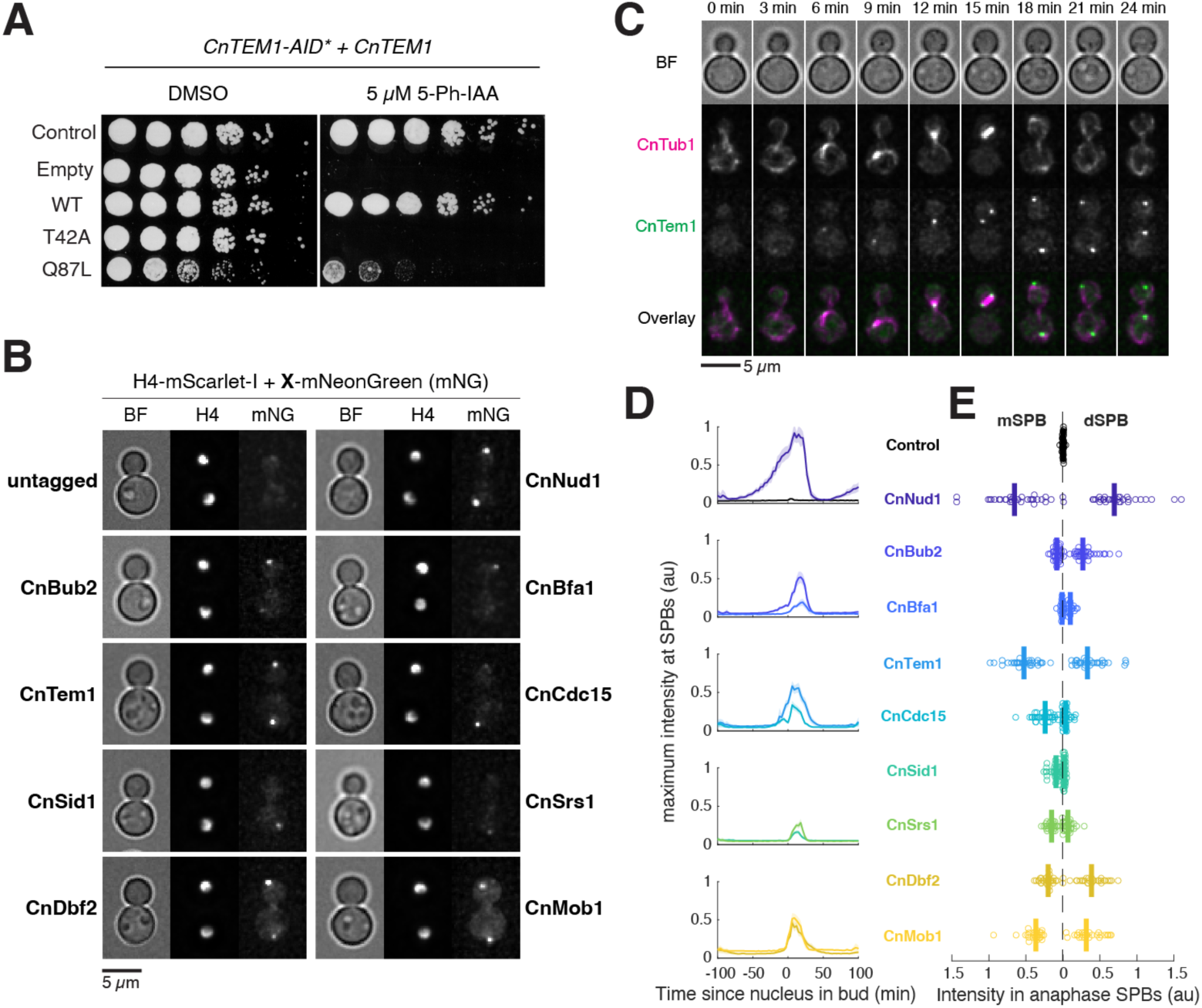
The *C. neoformans* MEN/SIN pathway signals via conserved mechanisms. (A) Complementation analysis for GTPase mutants of CnTem1. Serial dilutions of Cn278 (Control), Cn326 (Empty, *CnTEM1-AID*), Cn407 (WT, *CnTEM1-AID*), Cn407 (WT, *CnTEM1-AID* + *CnTEM1*), Cn408 (T42A, *CnTEM1-AID* + *Cntem1-T42A*), and Cn409 (Q87L, *CnTEM1-AID* + *Cntem1-Q87L*) were spotted onto YPD with DMSO or 5 µM 5-Ph-IAA and grown for 1.5 days at 30 °C. (B) Representative images of core MEN/SIN protein localization in anaphase cells. H4-mScarlet-I and C-terminal mNeonGreen fusions of CnNud1 (Cn106), CnTem1 (Cn125), CnCdc15 (Cn127), CnSid1 (Cn131), CnSrs1 (Cn139), CnDbf2 (Cn137), CnMob1 (Cn129), and untagged control (Cn84) were imaged on agarose pad made with SC media + 2% glucose at 30 °C. (C) Localization patterns of CnTem1 relative to tubulin during the cell cycle. Cn380 expressing CnTem1-mNeonGreen and mScarlet-I-CnTub1 was imaged on agarose pad made with SC media + 2% glucose at 30 °C. The pattern observed is consistent with SPB localization of CnTem1. (D) Localization dynamics of core MEN/SIN proteins during the cell cycle. Same strains and growth conditions as in (B). Solid lines represent the mean; shaded areas represent 95% confidence intervals. All components exhibit cell cycle dependent SPB localization. (E) Asymmetry of core MEN/SIN protein localization at the SPB in anaphase. Same data as (D) but maximum intensities for each protein in the mother cell (proxy for mSPB localization, left side of *x* axis) or daughter cell (dSPB, right side of *x* axis) were quantified for anaphase cells (*n* = 16, 31, 35, 19, 31, 37, 38, 28, 29, and 26 cells respectively for each strain).

Next, we characterized the cellular localization dynamics of each pathway component via endogenous fusion with the fluorescent protein mNeonGreen at the C-terminus. To determine the cell cycle stages, we also included a histone marker H4-mScarlet-I in these strains and performed time-lapse microscopy. We found that in anaphase cells all pathway components localized to puncta consistent with SPB localization (Fig. 2B, S2). Co-labeling with the spindle marker, α-tubulin CnTub1, confirmed the SPB localization (Fig. 2C). All pathway components displayed cell cycle dependent SPB localization including the scaffold protein CnNud1 (Fig. 2D) whereas its ortholog Nud1/Cdc11 is constitutively associated with the SPB as a core component throughout the cell cycle in *S. cerevisiae* and *S. pombe* respectively^31,47^. In basidiomycetes, the SPB is inactive as a MTOC in interphase and only becomes an active MTOC during mitosis to organize chromosome segregation^48,49^. Our observation revealed that the signaling function of SPB is also cell cycle regulated and precedes its activation as an MTOC by comparing the cell cycle localization dynamics of CnNud1 with the localization of γ-tubulin CnTub4 and α-tubulin CnTub1 (Fig. S3).

Finally, some MEN/SIN pathway components display asymmetric localization on the two anaphase SPBs in *S. cerevisiae*/*S. pombe*, where the GAP complex predominantly localizes to one SPB in both organisms (e.g., the SPB in the daughter cell or dSPB in *S. cerevisiae*).

Interestingly, the kinases Cdc7 and Sid1-Cdc14 in *S. pombe* localize to the opposite SPB from the GAP while in *S. cerevisiae* their counterpart Cdc15 localizes to both SPB. This asymmetry was shown to be important for correct signaling performance^44^. We found that in *C. neoformans* anaphase cells, the scaffold CnNud1, the GTPase CnTem1, and the kinase CnDbf2-CnMob1 localized to both SPBs while the GAP complex and the kinases CnCdc15 and CnSid1-CnSrs1 displayed localization asymmetry with a preference towards the dSPB and mSPB respectively (Fig. 2B, 2E).

### The MEN/SIN pathway in C. neoformans controls cytokinesis

To determine the function of the orthologous MEN/SIN genes we identified in *C. neoformans*, we characterized their cellular phenotype upon auxin-induced degradation. In *S. cerevisiae* MEN mutants arrest in anaphase generating large-budded cells with an elongated spindle as the MEN controls both mitotic exit and cytokinesis. In comparison, SIN mutants in *S. pombe* mainly have defects in cytokinesis/septation while the nuclear division cycle continues, leading to multi-nucleated cells. We first examined *CnTEM1-AID* cells expressing H4-mScarlet-I to track nuclear division and found that upon CnTem1 depletion cells did not arrest in anaphase. Rather, they continued to bud and entered the next round of nuclear division cycle (Fig. 3A). However, the budding pattern of these cells was quite distinct from control cells. In wild-type *C. neoformans* cells, the same bud site is used for many generations and there is a notable delay for the daughter cells to produce a bud after cell separation while the mother cells re-bud immediately (Fig. 3A Control). We found that instead of budding again from the mother cell at the same site, *CnTEM1-AID* cells frequently produced a new bud from the previous bud, generating cell chains (Fig. 3A, Chain), or budded from the bud neck (Fig. 3A, Shamrock), or another site on the mother cell (Fig. 3A, Mickey). The same phenotype was also observed for all the MEN/SIN genes (Fig. 3B, S4A) except the negative regulator GAP complex (CnBub2-CnBfa1), CnCdc15, and CnCdc14 which also failed to elicit a phenotype in growth assays.

**Figure 3.**
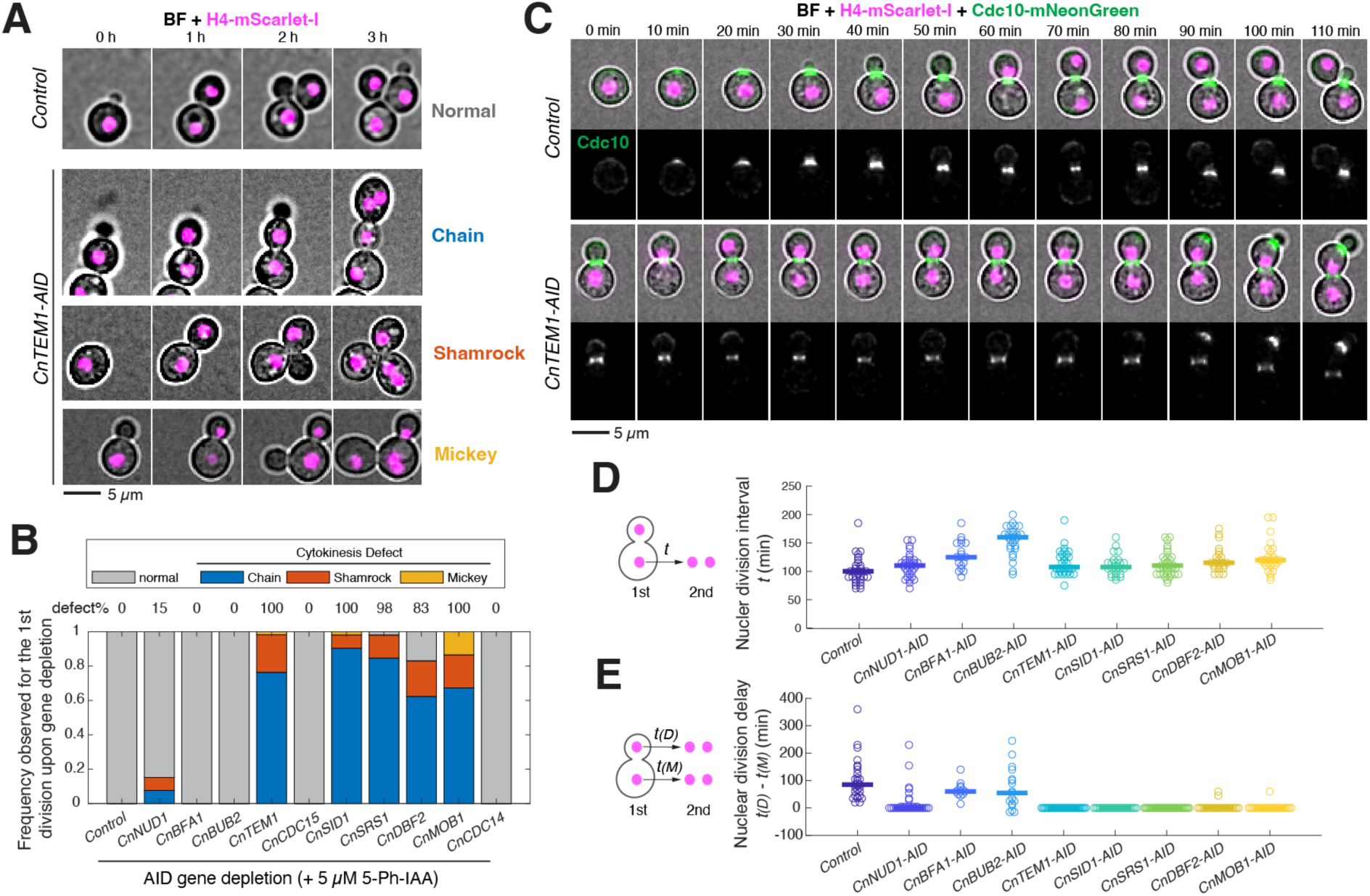
The MEN/SIN pathway regulates cytokinesis in *C. neoformans*. (A) Representative time-lapse images of cells with CnTem1 depleted expressing histone marker. Control (Cn278) and *CnTEM1-AID* (Cn325) cells expressing H4-mScarlet-I were imaged on agarose pad made with SC media + 2% glucose + 5 µM 5-Ph-IAA at 30 °C. Three different budding patterns following cytokinesis failure were labeled. (B) Frequency of cytokinesis defects observed for core MEN/SIN components gene depletion. *C. neoformans* strains with *CnNUD1-AID* (Cn41), *CnBFA1-AID* (Cn45), *CnBUB2-AID* (Cn53), *CnTEM1-AID* (Cn36), *CnCDC15-AID* (Cn37), *CnSID1-AID* (Cn39), *CnSRS1-AID* (Cn72), *CnDBF2-AID* (Cn44), *CnMOB1-AID* (Cn47), *CnCDC14-AID* (Cn49) and Control (Cn26) were imaged on SC media + 2% glucose + 5 µM 5-Ph-IAA at 30 °C. Only the phenotype of the first division upon 5-Ph-IAA exposure was scored for 50 cells per strain. (C) Representative time-lapse images of cells with CnTem1 depleted expressing histone and septin markers. Control (Cn347) and *CnTEM1-AID* (Cn344) cells expressing H4-mScarlet-I and Cdc10-mNeonGreen were imaged on agarose pad made with SC media + 2% glucose + 5 µM 5-Ph-IAA at 30 °C.(D-E) Nuclear division intervals (D) and delays (E) for cells with core MEN/SIN component depleted. *C. neoformans* strains with *CnNUD1-AID* (Cn341, *n* = 32), *CnBFA1-AID* (Cn337, *n* = 19), *CnBUB2-AID* (Cn339, *n* = 27), *CnTEM1-AID* (Cn326, *n* = 30), *CnSID1-AID* (Cn323, *n* = 25), *CnSRS1-AID* (Cn371, *n* = 36), *CnDBF2-AID* (Cn368, *n* = 23), *CnMOB1-AID* (Cn370, *n* = 27) and Control (Cn278, *n* = 32) were grown on SC media + 2% glucose + 5 µM 5-Ph-IAA at 30 °C and imaged every 5 min.

We hypothesized that this alternated budding pattern was due to a failure in cytokinesis as a similar budding phenotype has been previously reported in *S. cerevisiae* for cytokinesis defects^50^. Consistent with a cytokinesis failure, we observed multinucleation in the next nuclear cycle following the abnormal budding (see Fig. 3A, *CnTEM1-AID* 3h for examples) for all the mutants analyzed. To confirm and further characterize the cytokinesis defect, we fluorescently tracked the septin cytoskeleton that regulates cytokinesis by tagging the septin subunit CnCdc10 with mNeonGreen. In *S. cerevisiae*, septin assembles at the bud site and forms an hourglass-shaped collar during budding that helps recruit cytokinesis machinery^51^. During cytokinesis, the septin collar is restructured into two distinct rings and disassembles after cytokinesis^51^. Similar septin localization patterns have been observed in *C. neoformans*^52^ and were recapitulated in part by our CnCdc10 reporter (Fig. 3C Control). As mother cells typically re-bud immediately at the same site, we observed that the septin transitioned from ring to extended cone in the mother cell following cell separation while in the daughter cell the ring disappeared (Fig. 3C-Control 80-100 min). In contrast, the septin structure persisted at the bud neck in *CnTEM1-AID* cells after nuclear division and a second septin structure emerged associated with the new bud, (Fig. 3C *CnTEM1-AID*) consistent with defects in cytokinesis.

Furthermore, we followed the cells with MEN/SIN genes depleted to the next nuclear division and quantified the relative timing of nuclear divisions following cytokinesis failure. We found that in control cells on average it took about 100 minutes for the mother cell to undergo nuclear division following the previous nuclear division (Fig. 3D) and the daughter cell an extra 90 minutes (Fig. 3E) due to a G1 delay for cell growth (Fig. S5A). For all MEN/SIN genes with a cytokinesis defect when depleted (namely, CnNud1, CnTem1, CnSid1, CnSrs1, CnDbf2, and CnMob1), the delay for the daughter nucleus was eliminated as the two nuclei underwent mitosis synchronously (Fig. 3E). This observation confirms that the two nuclei share the same cytoplasm because of cytokinesis failure. Interestingly, it took on average about 10-20 minutes longer for these paired nuclei to divide compared to the mother nucleus in control cells, likely reflective of the difference in cytoplasm to nuclei ratio.

For the negative regulator GAP complex CnBub2 and CnBfa1 whose depletion did not result in cytokinesis failure, we observed a notable delay in the cell cycle progression where median nuclear division intervals increased from 100 min to 125 min for *CnBFA1-AID* and to 160 min for *CnBUB2-AID* cells (Fig. 3D). This delay is likely responsible for the slow growth phenotype we observed for these mutants on spotting assays (Fig. 1C). To identify which cell cycle stage contributed to the delay in GAP mutants, we tagged H4 with mNeonGreen which has a shorter maturation time (10 min) than mScarlet-I (36 min) and enabled us to monitor histone assembly during S phase with finer temporal resolution. Our cell cycle analysis indicated that there was a G2 delay in *Cnbub2Δ* cells generating large, budded cells where nuclei with replicated genome delayed entering mitosis by ∼50 minutes on average (Fig. S5A). As a result, we found that the daughter cells of this mutant were born with a larger size (Fig. S5B) and had a shorter G1 delay before entering mitosis (Fig. S5A, 3E). In rare occasions, the daughter cell entered mitosis before the mother cell (Fig. 3E, negative delays in nuclear division timing). We observed a similar phenotype in cells expressing the GTP-locked CnTem1-Q87L confirming that the mitotic delay is a result of hyperactivating CnTem1 (Fig. S4C).

Given the mitotic entry delay observed when hyperactivating CnTem1, we asked whether the MEN/SIN genes play a role in regulating mitosis in addition to cytokinesis in *C. neoformans*. To this end, we followed the spindle dynamics in *CnTEM1-AID* cells by expressing fluorescently labeled α-tubulin (mScarlet-I-CnTub1) together with H4-mNeonGreen for cell cycle timing analysis. We focused on the M phase which starts with nuclear/chromosome migration into the bud in *C. neoformans* and is followed by chromosome condensation (marked by a decrease in nuclear area and an increase in H4 intensity, Fig. S6) and spindle formation in the bud. Upon anaphase onset, the spindle elongates into the mother before breaking down at mitotic exit where chromosomes decondensed and the Tub1 signal returned to a less concentrated interphase microtubule network (Fig. S6). We compared the timing of mitotic entry and exit as defined by chromosome condensation (H4 intensity) and spindle formation (Tub1 intensity) and found no notable differences between the control and *CnTEM1-AID* cells in their first mitosis upon gene depletion (Fig. S6).

We concluded that all MEN/SIN orthologs in *C. neoformans* function in one pathway that regulates cytokinesis/septation.

### Dynein is required for nuclear migration during mitosis

Having established that the MEN/SIN pathway is conserved in *C. neoformans* and controls cytokinesis, we next examined whether spindle position regulates pathway activation, as in the ascomycetes budding yeasts *S. cerevisiae* and *C. albicans*, to coordinate cytokinesis with mitosis. To perturb spindle positioning in *C. neoformans*, we generated an auxin-inducible degron fusion of the heavy chain of cytoplasmic dynein, CnDyn1 (*CnDYN1-AID*), as *C. neoformans* lacks a Kar9 ortholog^53^ and Dyn1 has been implicated in nuclear migration in *C. neoformans*^54–56^. We found that unlike in *S. cerevisiae*, depleting CnDyn1 is lethal for colony formation in *C. neoformans* (Fig. 4A). This is consistent with the lack of a second Kar9-based pathway in *C. neoformans*, as in *S. cerevisiae* single *dyn1Δ* and *kar9Δ* mutants are viable but a double mutant is lethal^19^. We next monitored the spindle position in *CnDYN1-AID* cells by following fluorescently tagged H4 and Tub1 and demonstrated that nuclear migration to the bud upon mitotic entry was inhibited with CnDyn1 depletion and mitosis proceeded within the mother cell compartment (Fig. 4B). We concluded that *C. neoformans* relies solely on dynein to position the nucleus during mitosis and nuclear migration into the bud is not required to initiate mitosis.

**Figure 4.**
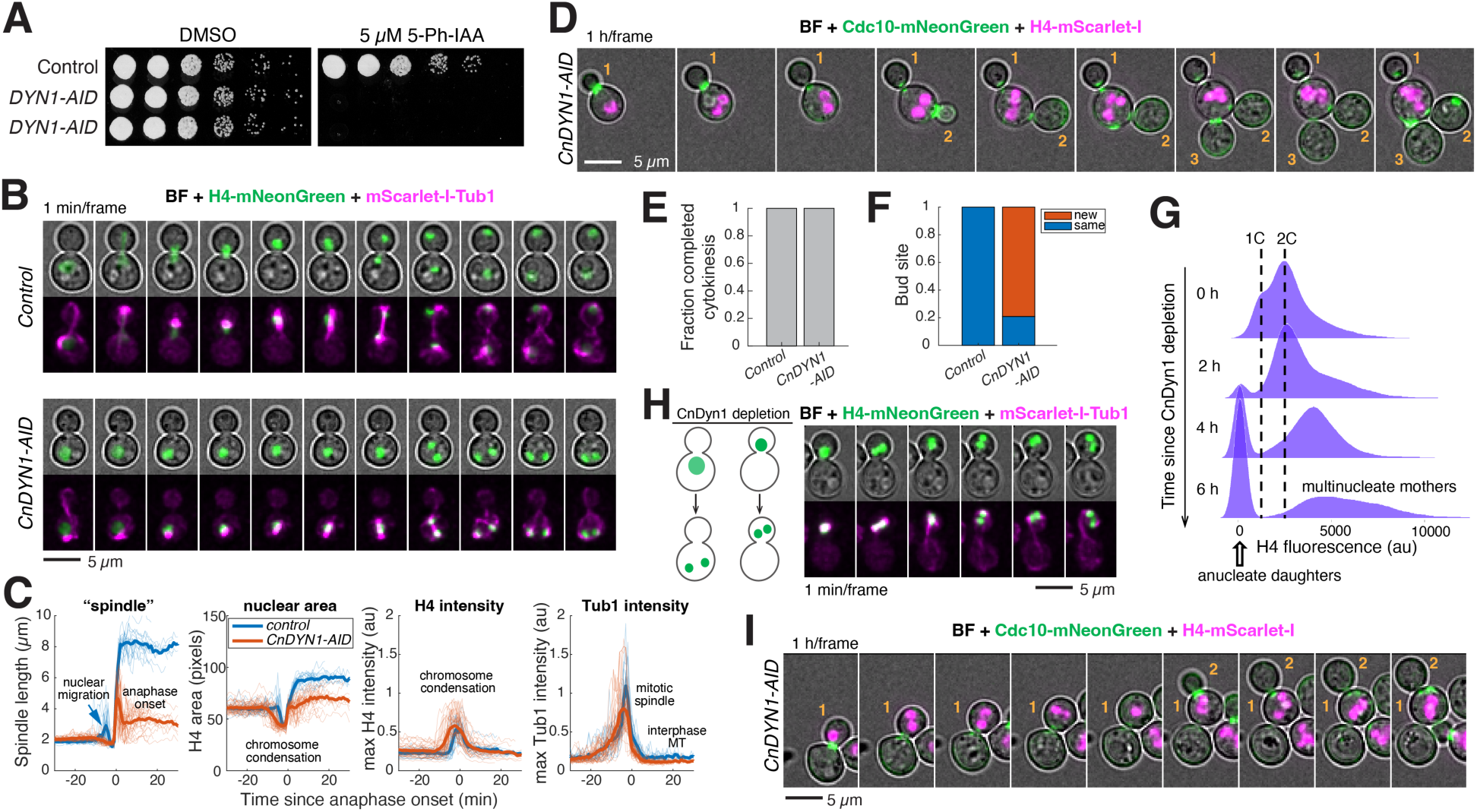
Spindle position does not regulate mitotic exit or cytokinesis in *C. neoformans*. (A) Colony formation analysis for *CnDYN1-AID*. Serial dilutions of Cn26 (Control), Cn57 (*CnDYN1-AID*), and Cn58 (*CnDYN1-AID*) were spotted onto YPD with DMSO or 5 µM 5-Ph-IAA and grown for 1.5 days at 30 °C. (B) Representative time-lapse images of cells with CnDyn1 depleted expressing fluorescent H4 and Tub1. Control (Cn364) and *CnDYN1-AID* (Cn349) cells expressing H4-mNeonGreen and mScarlet-I-Tub1 were grown on agarose pad made with SC media + 2% glucose + 5 µM 5-Ph-IAA at 30 °C. (C) Comparison of mitosis timing for *CnDYN1-AID* cells. Same cells and growth conditions as in (B). Spindle length (estimated by the long axis of H4 signal), nuclear area (area of H4 signal), H4 intensity (maximum intensity of H4 in the cell), and Tub1 intensity (maximum intensity of Tub1 in the cell) were quantified for each cell and the single cell traces (thin lines) were aligned based on the time of anaphase onset (spindle length > 3 µm) and averaged (thick lines). 18 cells for Cn364 and 42 cells for Cn349 were analyzed. (D) Representative time-lapse images of cells with CnDyn1 depleted expressing fluorescent H4 and septin. *CnDYN1-AID* cells expressing H4-mScarlet-I and Cdc10-mNeonGreen (Cn389) were imaged as in (B). Numbers label the anucleate buds produced upon CnDyn1 depletion. (E-F) Frequency of cytokinesis (E) and bud site switching (F) for Control (Cn26) and *CnDYN1-AID* cells (Cn58). Same growth conditions as in (B), 24 cells were scored for each strain. (G) Ploidy of *CnDYN1-AID* cells over time upon CnDyn1 depletion. Cn181 was grown in YPD to log phase at 30 °C and 5 µM 5-Ph-IAA was added at 0 hours. (H-I) Representative time-lapse images of cells with CnDyn1 depleted after nuclear migration. Cn349 (H) and Cn389 (I) were grown and imaged as in (B) and (D). Bud #1 retained both nuclei as CnDyn1 depletion occurred after nuclear migration and continued to make anucleate bud (#2).

### Spindle position does not regulate mitotic exit or cytokinesis in C. neoformans

To examine whether *C. neoformans* cells monitor spindle position and regulate cell cycle progression accordingly, we followed the fate of *CnDYN1-AID* cells where nuclear division occurred within the mother cell upon depletion. We hypothesized that, given the spatiotemporal constraint of division by budding, spindle position could either regulate cytokinesis via the MEN/SIN pathway like in ascomycetes yeasts or regulate mitotic exit through an alternative mechanism in *C. neoformans* to safeguard the fidelity of mitosis. To our surprise, we found that neither was inhibited upon failure of nuclear migration to the bud as both mitotic exit (monitored with chromosome decondensation via H4 intensity and spindle disassembly via Tub1 intensity, Fig. 4C) and cytokinesis (monitored with brightfield and septin marker Cdc10, Fig. S7) ensued with similar kinetics relative to anaphase onset as control cells.

As both nuclei were retained in the mother cell, this division resulted in a bi-nucleated mother and anucleate daughter. Following *CnDYN1-AID* cells for an extended period revealed that, impressively, the bi-nucleated mother cell went on to the next cell cycle and budded off anucleate daughters for several generations (Fig. 4D) without cytokinesis failure (Fig. 4E), in stark contrast to the MEN-AID mutant phenotype (Fig. 3). This is further validated by flow cytometry analysis of *CnDYN1-AID* cells where anucleate and multinucleate cell populations accumulated over time upon gene depletion (Fig. 4G). Interestingly, we noticed that unlike control cells which reused the same bud site for successive generations, *CnDYN1-AID* cells frequently switched the bud site (Fig. 4F).

Our observation that failure of nuclear migration to the bud does not arrest cell cycle progression or block cytokinesis suggests that there might not be a SPoC in *C. neoformans*. Since the mitotic spindle is assembled in the bud in *C. neoformans*, we reasoned that perhaps the SPoC, if exists, could monitor the return of a nucleus (a set of chromosomes) to the mother compartment instead. Interestingly, we found that a small fraction of the *CnDYN1-AID* cells, where gene depletion occurred after the nucleus had already migrated into the bud, underwent nuclear division within the bud without returning a nucleus to the mother. Tracking the fate of this subpopulation revealed that cell cycle progression was not inhibited by anucleate mother either as these cells continued with exit from mitosis and cytokinesis (Fig. 4H-I). Furthermore, the bi-nucleated daughters proceeded to bud (Fig. 4I) albeit with a slight delay compared to control cells, likely due to the increased nuclear to cytoplasmic ratio at birth.

We concluded that correct spindle positioning is not required for *C. neoformans* MEN/SIN pathway activation to drive cytokinesis or for cell cycle progression to continue. In essence, there does not appear to be a SPoC in this organism despite having the same division plane constraint of budding as ascomycetous budding yeasts. Furthermore, our results with *CnDYN1-AID* suggest that dynein motor is required for both nuclear migration to the bud upon mitotic entry as well as the return of a nucleus to the mother cell in anaphase.

### Failure in nuclear division does not block cytokinesis in C. neoformans

Given the absence of a SPoC in *C. neoformans*, we asked whether there is an alternative surveillance mechanism that regulates cytokinesis in response to defects in nuclear division or chromosome segregation. To this end, we perturbed the mitotic spindle by treating cells with the microtubule-depolymerizing agent nocodazole. To avoid metaphase arrest, we introduced *mad2Δ* to bypass the spindle assembly checkpoint (SAC)^15^, which inhibits metaphase to anaphase transition in response to defects in kinetochore to spindle microtubule attachments^57^. As reported previously^15^, nocodazole treatment arrested control *C. neoformans* cells in metaphase with condensed chromosomes in the mother cell as marked by H4-mNeonGreen (Fig. 5A WT). In contrast, *mad2Δ* cells exited mitosis (as marked by chromosome decondensation) and rebudded^15^ with similar kinetics as untreated cells (Fig. 5A *mad2Δ*) despite not having undergone nuclear division (chromosome segregation). Importantly, we found that these cells also completed cytokinesis without a delay compared to untreated cells (Fig. 5B) giving rise to anucleate daughters and diploidized mothers. The diploidized mothers immediately entered the next cell cycle and continued to bud off anucleate daughters while doubling its ploidy each cycle as a result (Fig. 5C). These observations revealed that there does not appear to be any surveillance mechanism to block cytokinesis upon defective mitosis in *C. neoformans*.

**Figure 5.**
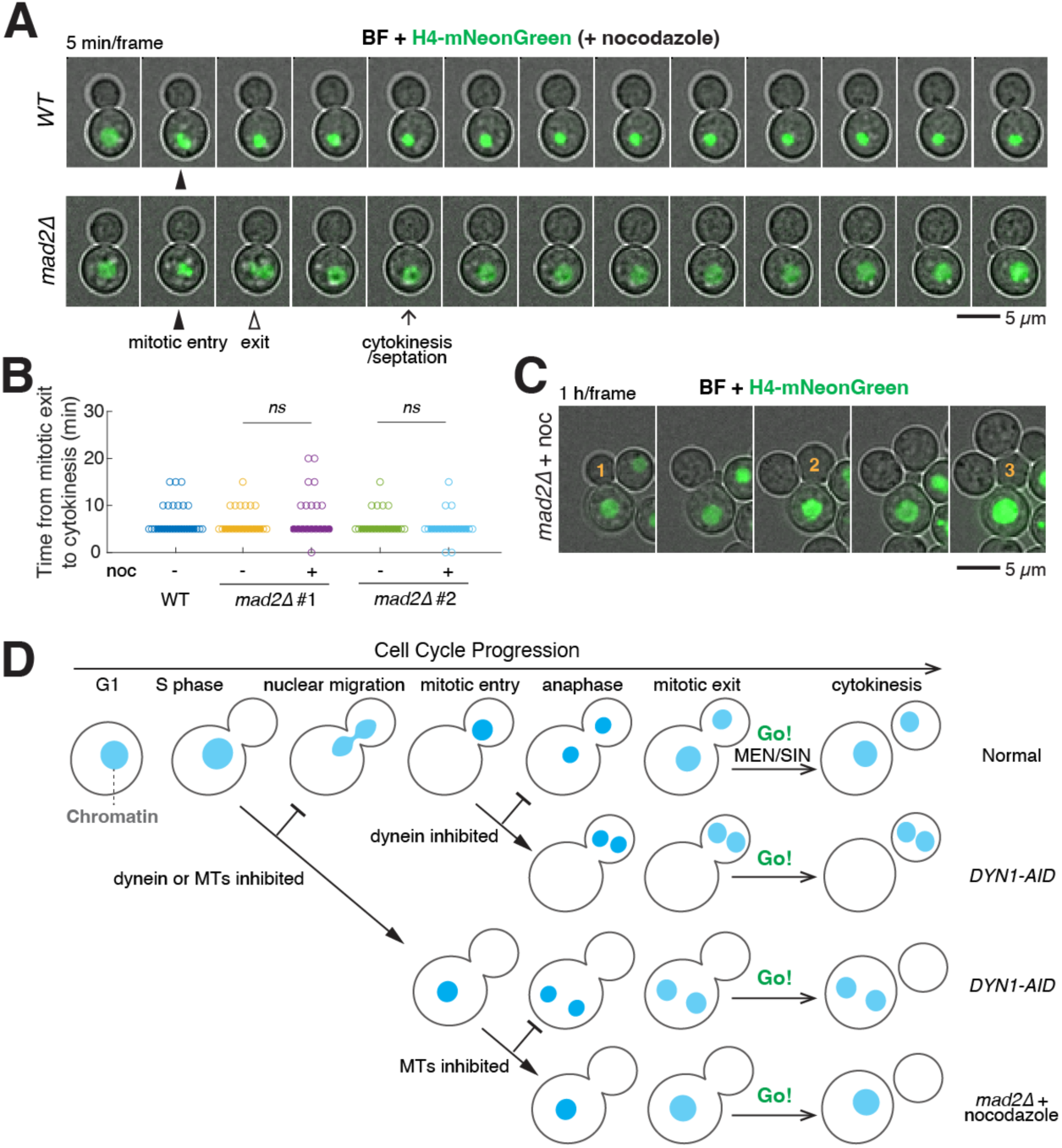
No checkpoint to delay cytokinesis upon mitotic defects in *C. neoformans*. (A) Representative time lapse images of fluorescent H4 expressing cells treated with nocodazole. WT (Cn189) and *mad2Δ* (Cn435) cells expressing H4-mNeonGreen were grown on agarose pad made with SC media + 2% glucose + 5 µg/ml nocodazole and imaged every 5 minutes. Mitotic entry and exit were marked based on H4 signal (chromosome condensation) and cytokinesis was scored based on the bright field image (appearance of dark line at bud neck). (B) Comparison of cytokinesis timing relative to mitotic exit for nocodazole treated cells. Same strains and conditions as in (A). Each dot denotes a cell (*n* = 29, 22, 23, 21, 21 cells respectively) and solid lines denote the medians. *ns*, not significant (*p* = 0.3 and 0.2 respectively) by two-sided Wilcoxon rank sum test. (C) Successive rounds of cell division for nocodazole treated *mad2Δ* cells. Cn435 was grown as in (A) and imaged every 5 minutes for 8 hours. Numbers indicate the anucleate buds generated. (D) Model for *C. neoformans* mitosis and cytokinesis. In unperturbed cell cycle, mitosis starts with dynein-mediated nuclear migration into the bud which is followed by mitotic entry (marked by chromosome condensation and spindle formation) in the bud. Upon anaphase onset, chromosome segregation proceeds across the bud neck with the help of dynein and is followed by mitotic exit (marked by chromosome decondensation and spindle disassembly). Cytokinesis is controlled by the conserved MEN/SIN pathway and occurs shortly after mitotic exit. When nuclear movement or chromosome segregation was inhibited with *DYN1-AID* or nocodazole treatment in *mad2Δ* cells respectively, cytokinesis proceeded as normal generating anucleate and multinucleate or polyploid progeny.

## DISCUSSION

In this study, we investigated the coordination between cytokinesis and mitosis in the fungal pathogen *C. neoformans*. We found that the MEN/SIN pathway, a GTPase-kinase signaling cascade that regulates cytokinesis in ascomycetes, is conserved in the basidiomycetous budding yeast *C. neoformans* and controls cytokinesis. However, unlike ascomycetous yeasts, the *C. neoformans* MEN/SIN pathway is not regulated by spindle position, rendering this organism devoid of a checkpoint that delays cytokinesis upon defects in mitosis.

The MEN/SIN pathway is ancient in eukaryotes with varying functional divergence among different lineages. In animals the homologous pathway is known as Hippo pathway and regulates tissue proliferation and organ size^58^. In ascomycetous fungi, the pathway mainly regulates cytokinesis, and it was previously unclear whether this represents the ancestral pathway function in fungi as a different function (nuclear envelope breakdown) was described for the only non-ascomycete fungus (*Ustilago maydis*) studied so far. Our study in *C. neoformans*, a basidiomycetous budding yeast, suggests that the cytokinesis role of the MEN/SIN pathway is conserved beyond Ascomycota and is likely ancestral in the common ancestor of Ascomycota and Basidiomycota. Future studies in additional species both in Basidiomycota as well as other fungal lineages will help shed light on the ancestral role and evolution of the MEN/SIN pathway in fungi.

Despite the overall conservation of the pathway in fungi, we observed distinct variations among organisms. Most notably, the MEN in *S. cerevisiae* has modified both pathway composition (lost Sid1) and function (acquired a major role in regulating mitotic exit). We propose that the latter likely resulted from a functional rewiring of the downstream target, the phosphatase Cdc14, which is essential in *S. cerevisiae* regulating mitotic exit^59^ and nonessential in *S. pombe*^60,61^, *C. albicans*^62^, and *C. neoformans* in our study. Another major difference revealed by our study is the essentiality of the GAP complex or the consequences of hyperactivating the pathway at the GTPase step. In *S. cerevisiae*, constitutively activating the GTPase Tem1 bypasses the SPoC but has little impact on the cell cycle (slight acceleration of mitotic exit)^42^. In contrast, constitutively active GTPase Spg1 is lethal in *S. pombe* resulting in multiseptated cells^46^. In *C. neoformans*, constitutively activating the GTPase either with a GAP mutant (*Cnbub2Δ*) or a GTP-locked mutant of CnTem1 leads to cell cycle delays particularly for mitotic entry, which is distinct from both model yeasts. This difference among organisms likely reflects the variations in how the pathway is wired to the rest of the cell cycle and whether additional layers of regulations exist downstream of the GTPase.

We do not yet know what the upstream signals are for the *C. neoformans* MEN/SIN pathway. In *S. cerevisiae* and *S. pombe*, both temporal cues (cell cycle progression) and spatial cues (spindle position or spindle elongation) regulate pathway activation to ensure cytokinesis only occurs after proper mitosis. Given the lack of regulation from any spatial cues of mitosis revealed by our study, we propose that *C. neoformans* MEN/SIN pathway is mainly regulated by cell cycle progression. In both the MEN and SIN, polo-like kinase (Cdc5/Plo1), which is only expressed and active in mitosis^63^, is a key activator of the pathway^64–66^ and high mitotic cyclin-dependent kinase (CDK) activity inhibits the pathway^31,67^. This combination limits full pathway activation to anaphase when CDK activity has dropped from the metaphase peak and Cdc5/Plo1 is active.

By perturbing spindle positioning, our analysis also revealed new insights for mitosis in *C. neoformans*. Unlike the well characterized model budding yeasts in the Ascomycota, mitosis in *C. neoformans* starts with nuclear/chromatin migration into the bud where the mitotic spindle is formed and chromosomes segregate from the bud to the mother cell. It was not fully understood how this noncanonical chromosome segregation/nuclear division is achieved and whether nuclear migration into the bud is required for mitotic entry. By depleting CnDyn1, we showed that the microtubule minus-end motor dynein is required for both nuclear migration to the bud upon mitotic entry and chromosome segregation back to the mother cell in anaphase. Furthermore, our results demonstrated that mitosis could proceed in the mother cell in the absence of nuclear migration albeit with a slight delay in anaphase onset relative to mitotic entry (Fig. 4C, note the shifted onset of chromosome condensation as shown in nuclear area and H4 intensity). This delay could be a result of defects/delays in nuclear envelope breakdown which has been linked to nuclear migration in another basidiomycete yeast *Ustilago maydis*^37^. Nevertheless, our analysis suggests that instead of being a prerequisite for mitosis, nuclear migration is independent from mitosis. One possibility is that dynein-mediated nuclear migration could be regulated by mitotic CDK which also drives mitotic entry. This would provide a built-in mechanism to temporally coordinate these two events in a normal cell cycle.

To maintain genome stability particularly in mononucleated cells, cytokinesis needs to occur after the successful completion of mitosis, and the site of cytokinesis (division plane) and spindle position/orientation must be tightly coordinated. A lack of coordination could have detrimental consequences for the progeny and as a result surveillance mechanisms (i.e., checkpoints) have evolved in many organisms to delay cytokinesis when errors in mitosis occur, such as upon spindle mispositioning or damage. We showed here that *C. neoformans* does not appear to have such a checkpoint via two orthogonal approaches. By genetically depleting CnDyn1, we inhibited proper spindle positioning during mitosis and generated cells with both nuclei in the mother compartment or the bud. By treating cells with the microtubule poison nocodazole, nuclear movement as well as spindle formation were disrupted. In both cases, we found little delays in cytokinesis relative to anaphase onset or mitotic exit resulting in the accumulation of anucleate and multinucleate (*CnDYN1-AID*) or polyploid (nocodazole) progeny (Fig. 5D).

This is in stark contrast to both model yeasts *S. cerevisiae* and *S. pombe*. In *S. cerevisiae*, SPoC arrests cells in anaphase with cytokinesis blocked upon spindle mispositioning as the MEN regulates both mitotic exit and cytokinesis (Fig. 1A). Constitutive activation of the MEN GTPase (e.g., with the GAP mutant *bub2Δ*) bypasses the SPoC and produces anucleate daughters and binucleate mothers when anaphase spindle remains in the mother cell^23–25^. Moreover, cytokinesis does not occur in nocodazole treated *S. cerevisiae mad2Δ* cells^68^ despite them proceeding to the next cell cycle (re-budding and DNA re-replication) after a notable delay. Only when combined with *bub2Δ* would the cells proceed with mitotic exit and cytokinesis without a delay^69,70^. Similarly, *S. pombe* cells delay cytokinesis upon spindle stress^71^ and *mad2Δ* cells treated with microtubule-depolymerizing drug methyl-2-benzimidazole-carbamate (MBC) continue to cycle but fail to cytokinesis^36^. Hyperactivating the SIN bypassed this cytokinesis block. Finally, in *C. albicans*, a Bub2-dependent spindle position checkpoint delays mitotic exit and cytokinesis upon spindle mispositioning in the yeast form^35^. In essence, *C. neoformans* cells behave like *bub2Δ S. cerevisiae* or *C. albicans* cells as being SPoC deficient.

Our results with nocodazole treatment also demonstrated that due to the lack of SPoC *C. neoformans* cells could face a higher risk of genome instability in the presence of microtubule poison. What might be the benefit of not having a SPoC? As a human pathogen, *C. neoformans* has a fascinating life cycle in the host where the typical small haploid yeast undergoes an unusual morphogenic switch to form giant, highly polyploid “titan cells” with up to a 1000-fold increase in cell volume^3,4^ to escape phagocytosis by host immune cells^72^. It has been proposed that titan cells form through endoreduplication^7^, but the exact mechanism remains elusive. One possible model for endoreduplication is to skip mitosis while continuing the budding and DNA replication cycles by downregulating/suppressing mitotic cyclins. In this model, the lack of a SPoC would be beneficial to facilitate timely cytokinesis in the absence of nuclear migration and chromosome segregation during polyploidization, like the nocodazole treated cells in our study (Fig. 5C). Future comparative studies on nonpathogenic relatives of *C. neoformans* could help shed light on this possibility and the evolution of SPoC in fungi.

## ACKNOWLEDGEMENTS

We would like to thank Hiten Madhani and Kevin Hardwick for sharing strains and plasmids, Andrew Murray and Kevin Hardwick for suggestions on the manuscript. This work was supported by an NIH grant (DP2AI192733) to XZ.

## AUTHOR CONTRIBUTIONS

Conceptualization, T.K.W. and X.Z.; investigation, T.K.W., J.M., A.S., and X.Z.; resources, E.W.J.W.; formal analysis, T.K.W. and X.Z.; funding acquisition, X.Z.; supervision, X.Z.; visualization, T.K.W. and X.Z.; writing – original draft, T.K.W. and X.Z.; writing – review & editing, all authors.

## MATERIALS AND METHODS

### Growth and media conditions

All *Cryptococcus neoformans* strains used in this study are derivatives of KN99α and are listed in Table S1. For each strain generated, at least two independent transformants were isolated and characterized. Strains were routinely cultured in standard YEP media (1% yeast extract, 2% peptone) with 2% D-glucose (YPD), or in standard Synthetic Complete media (SC) with 2% D-glucose. Cells were cultured at 30 °C unless noted otherwise.

### Construction of plasmids

All plasmids used in this study are listed in Table S2. Ectopic Safe Haven integration plasmid^73^ for the OsTIR ligase^41^ and tagging plasmids for fluorescent fusions^74^ or AID fusions^41^ were constructed via NEB HiFi Gibson assembly to swap the resistance markers or fluorescent proteins. A Cryptococcus-optimized mScarlet-I coding sequence was synthesized. The mScarlet-I protein sequence^75^ was taken from the FPbase fluorescent protein database^76^. Codon usage for mScarlet-I was randomly generated weighted by the codon frequency table for the most translated 5% of CDS in H99^77^. The N-terminal fluorescently tagged CnTub1 plasmid was generated by amplifying the promoter, gene body, and terminator of CnTub1 via PCR from *C. neoformans* genomic DNA and cloning it into a Safe Haven 3 integration plasmid. A subsequent cloning step inserted mScarlet-I or mNeonGreen and a linker N-terminally of the CnTub1 start codon. A similar strategy of PCR from *C. neoformans* genomic DNA was utilized to clone the plasmids expressing wild type, GTP-locked, and GDP-locked versions of CnTem1 in the Safe Haven 3 integration vector, utilizing Q5 directed mutagenesis for single amino acid mutations.

### Construction of C. neoformans strains

The parent strain utilized in all strain engineering constitutively expressed a codon optimized Cas9^74^ (Cn23/ CM2049). To enable high-efficiency Cas9-mediated strain engineering the non-homologous end joining protein *CnKU80* was disrupted^78^ with the *amdS2* Blaster^73^ to generate Cn81 which was used for all fluorescent tagging strain construction. CM2476 (Cn26) expressing OsTIR ligase was used as the base strain for the AID C-terminal tagging to generate Cn35-72. A marker-less and reporter-less OsTIR ligase for the AID system was generated (p3171) and integrated at the Safe Haven 1 genomic site leveraging the *amdS2* Blaster to make Cn122. Cn122 was the base strain for all the rest of AID degron C-terminal tagging strains.

Cas9-mediated genetic engineering of *C. neoformans* strains was carried out as described previously^74^ with the following modifications. The repair templates were generated by PCR with ∼40 base pairs flanking homology for tagging the gene of interest with fluorescent proteins (p3160, p3192, p3195, or p3292) or AID (p3093) C-terminally. The sgRNAs containing necessary expression machinery were synthesized with TwistBio (sgRNA sequences used in this study can be found in Table S3). Specifically, electrocompetent cells were mixed with transformation DNA (> 2 µg of linearized Safe Haven plasmid DNA, or > 1 µg 40-base pair homology tagging construct and > 100 ng of appropriate guide RNA) and transferred to a pre-chilled 0.2 cm BioRad electroporation cuvette. Electroporation was carried out with a BioRad MicroPulser set to Sc2. Post-electroporation cells were recovered for 2 hours in liquid YPD at 30°C in a rotator and then plated onto solid YPD media supplemented with the appropriate selective drugs (125 ng/µL nourseothricin or 50 ng/µL G418). Correct engineering of colonies was verified via PCR screening and appropriate phenotype checking (e.g. fluorescence). PCR screening was carried out using primers binding upstream and downstream of the repair template homology. A lithium acetate (LiOAc)-SDS based genomic DNA extraction protocol^79^ was adapted for *C. neoformans* to isolate template DNA for checking PCRs.

For strains engineered with the recyclable *amdS2* Blaster system, cells were recovered for 1.5 hours in liquid YPD at 30°C in a rotator before plating cells on YNB media (0.45% yeast nitrogen base w/o amino acids and ammonium sulfate) containing 2% glucose, and 5 mM acetamide. To select for the popout event (self-excising of *amdS2* marker via the flanking direct repeats), cells were first patched on YPD plate and grown for 1-2 days at 30°C and a 5ml YPD culture was inoculated and grown overnight. ∼1 million cells were plated on YNB containing 2% glucose, 10 mM ammonium sulfate, and 10 mM fluoroacetamide. After 2-3 days colonies were checked for loss of *amdS2* cassette by PCR and growth on acetamide and fluoroacetamide.

### Identification of orthologs

Orthologs of MEN/SIN genes were identified through reciprocal BLAST between the *C. neoformans* H99 genome and *S. cerevisiae* S288C genome or *S. pombe* 972h-genome (for Sid1 and Cdc14).

#### Microscopy and image analysis

For live-cell microscopy, cells were imaged on agarose pads (2% agarose in SC medium + 2% glucose, unless otherwise noted) affixed to a glass slide and covered with a coverslip. Imaging was performed on a Nikon Ti2E with a SPECTRA III light source, controlled by NIS Elements software. A 60x Plan APO 1.42NA objective and ORCA-FUSIONBT SCMOS camera were used for image acquisition. For each time point, 7 *z* sections with 1 µm spacing were collected for each channel and were deconvolved. Maximum projections of the deconvolved *z* stack were used for fluorescence quantification.

Image analysis was performed with custom scripts in MATLAB. First, yeast cells were segmented and tracked through time using the bright-field image stacks. Next, fluorescence images of histone H4 were segmented and tracked based on cell segmentation. Appearance of H4 signal in a cell during the acquisition period was used to identify buds (daughter cells) and cell division events. Tracking of the H4 signal that migrated into the buds were used to identify the corresponding mother cells. Finally, for each division event identified, fluorescence intensities (total/sum and maximum) of H4 and additional markers were quantified for both the cellular compartments and nucleus.

Single cell traces were aligned based on the timing of nuclear migration or anaphase onset (estimated spindle length as defined by the long axis of H4 signal > 3µm) and averaged. 95% confidence intervals were calculated as 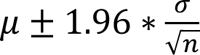, where *μ* and *σ* denote the mean and standard deviation respectively and *n* is the number of cells measured.

Manual scoring of nuclear division timing and cytokinesis timing and defects for strains was carried out as follows. Single cells were followed through time-lapse microscopy image series where the time frame of first nuclear division, and second nuclear divisions in mother and daughter cells were recorded. Nuclear division time frame was decided by the first time frame that showed two separate nuclei. Cytokinesis fates for single cells were scored based on whether the mother cell underwent cytokinesis (Normal) before budding again, or whether cytokinesis failed with subsequent budding at daughter cell (Chain), from mother-bud neck junction (Shamrock), or from the mother again (Mickey). Cytokinesis/septation timing was scored using bright field images based on the appearance of a dark line at bud-neck junction between mother and daughter cells.

#### Spotting assays

For spotting assays, *C. neoformans* strains were grown in YPD overnight and diluted to OD 0.2 in YPD and grown for at least one doubling at 30°C. 5-fold serial dilutions were made for each culture starting from OD 0.2 and 4 µl were spotted onto YPD plates containing dimethyl sulfoxide (DMSO) alone or 5-phenyl-indole-3-acetic acid (5-Ph-IAA, SML3574 from Millipore Sigma) which induces degradation of auxin tagged proteins. Stock solutions of 5Ph-IAA was made at 10 mM in DMSO.

#### Immunoblot analysis

*C. neoformans* log phase cells grown in YPD were harvested and treated with 5% TCA at 4°C overnight. TCA treated cells were pelleted and washed with 1M Tris (unbuffered) and resuspended in lysis buffer (10 mM Tris, 1 mM EDTA, 2.75 mM DTT, pH = 8). Approximately 100 µL of glass beads were added to each sample, and then samples were run on a FastPrep-24 for 3 cycles of speed 6.5 for 60 seconds with 5 minutes rest between cycles to prevent sample overheating. 30 µL of 4x sodium dodecyl sulfate (SDS) sample buffer was added and samples subsequently boiled at 95°C for 5 minutes. Lysates were clarified by centrifugation and resolved with NuPAGE 4-12% Bis-Tris protein gel (Thermo Fisher Scientific) prior to transfer onto nitrocellulose membranes. FLAG tag was detected using an anti-FLAG M2 antibody (F3165, Millipore Sigma) at a 1:1000 dilution. DyLight 800 conjugated anti-mouse secondary antibodies (5257S, Cell Signaling) were used at a 1:10,000 dilution. Blots were imaged using the ChemiDoc MP imager system (BioRad).

#### Flow Cytometry Analysis

Ploidy analysis was performed with H4-mNeonGreen. Log phase cells treated with DMSO or 5 µM of 5-Ph-IAA were harvested and fixed with 3.7% formaldehyde for 15 minutes. Samples were then resuspended in 100 mM potassium phosphatase, 1.2 M sorbitol solution with 1% triton for 5 minutes. Final resuspension was in 100 mM potassium phosphatase 1.2 M sorbitol solution. Cells were then sonicated at 20% amplitude for two 5 second on pulses with an intervening 5 second off. At least 10,000 events were collected via a CyTek Aurora flow cytometry using the BG-3 channel for collection. Analysis was performed via a custom RStudio pipeline in the lab.

## Notes

### Competing Interest Statement

The authors have declared no competing interest.

