## Supplementary Figures for "Absence of a spindle position checkpoint in the fungal pathogen *Cryptococcus neoformans*"

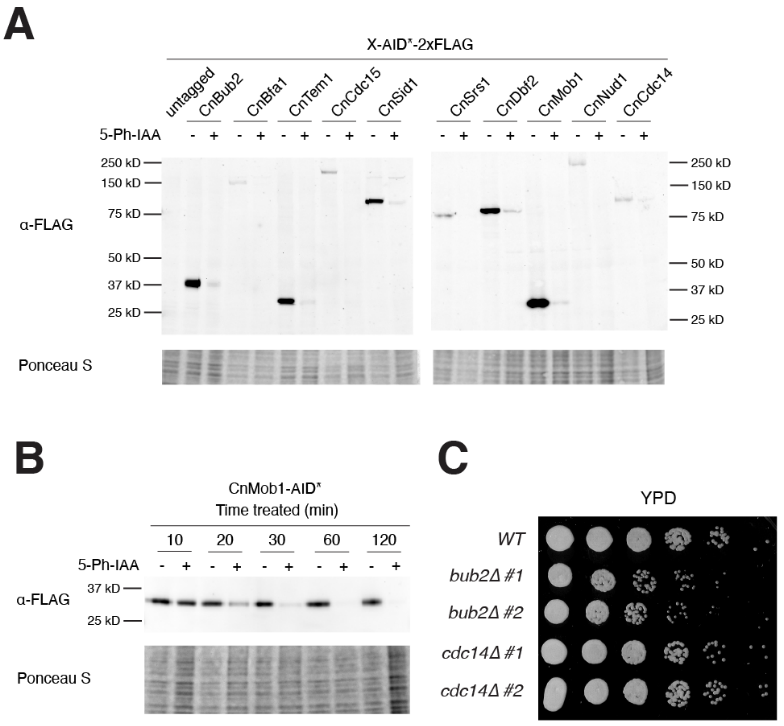


**Figure S1. Auxin-inducible degradation of core MEN/SIN components in *C. neoformans*.**

(A) Western blot analysis of the auxin-inducible degradation for the core MEN/SIN components. Cn26 (untagged), Cn53 (*CnBUB2-AID*), Cn45 (*CnBFA1-AID*), Cn35 (*CnTEM1-AID*), Cn37 (*CnCDC15-AID*), Cn39 (*CnSID1-AID*), Cn72 (*CnSRS1-AID*), Cn43 (*CnDBF2-AID*), Cn47 (*CnMOB1-AID*), Cn41 (*CnNUD1-AID*), and Cn49 (*CnCDC14-AID*) were grown in YPD to log phase and either DMSO or 5 μM of 5-Ph-IAA was added to induce degradation. Cells were harvested after 2 hours and processed for Western blot analysis. Anti-FLAG was used to probe for AID tagged proteins (AID*-2xFLAG). Total protein stained with Ponceau S was used as loading control.

(B) Kinetics of auxin-inducible degradation. Cn47 was grown in YPD to log phase and either DMSO or 5 μM of 5-Ph-IAA was added to induce degradation. Cells were harvested after the indicated time and processed for Western blot analysis. The degradation was complete by 1 hour.

(C) Growth analysis for *bub2*Δ and *cdc14*Δ. Serial dilutions of Cn81 (WT), Cn272 (*bub2*Δ), Cn273 (*bub2*Δ), Cn274 (*cdc14*Δ), and Cn275 (*cdc14*Δ) were spotted onto YPD and grown for 1.5 days at 30 ºC.

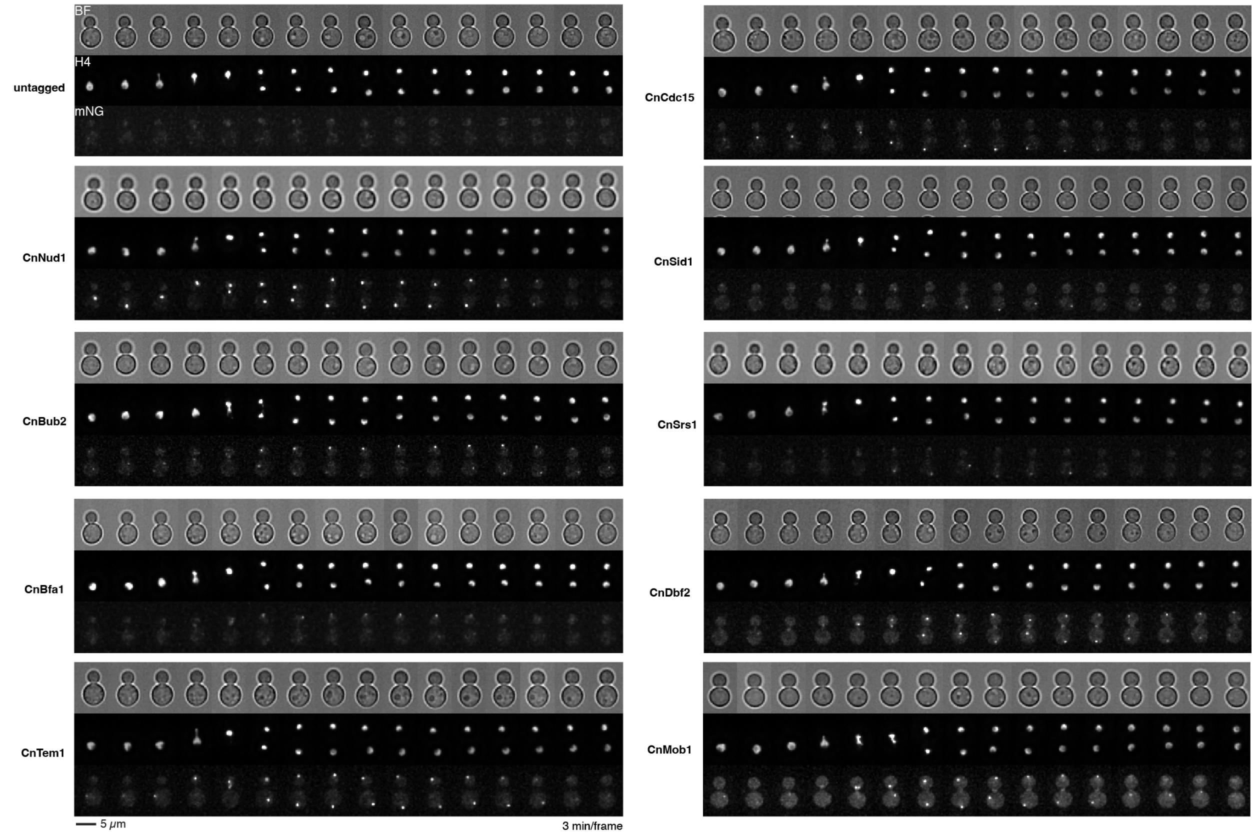


**Figure S2. Localization of core MEN/SIN proteins during the cell cycle in *C. neoformans*.**

Same strains and growth conditions as in Figure 2B. Cells were imaged every 3 min.

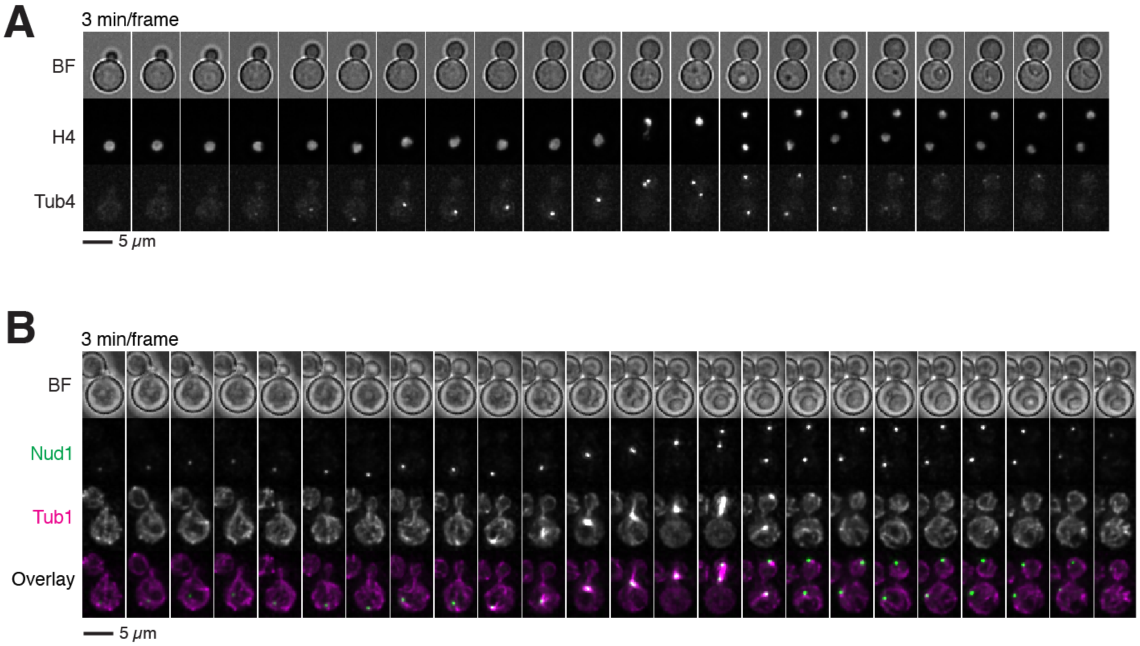


**Figure S3. CnNud1 localization to the SPB is cell cycle dependent**

(A) Localization pattern of γ-tubulin CnTub4 during the cell cycle. Cn107 expressing CnTub4-mNeonGreen and H4-mScarlet-I was imaged on agarose pad made with SC media + 2% glucose at room temperature.

(B) Localization pattern of CnNud1 relative to α-tubulin during the cell cycle. Cn332 expressing CnNud1-mNeonGreen and mScarlet-I-CnTub1 was imaged on agarose pad made with SC media + 2% glucose at 30 ºC. Note that CnNud1 started to localize to the SPB (bright puncta) before Tub1 when the bud was still small.

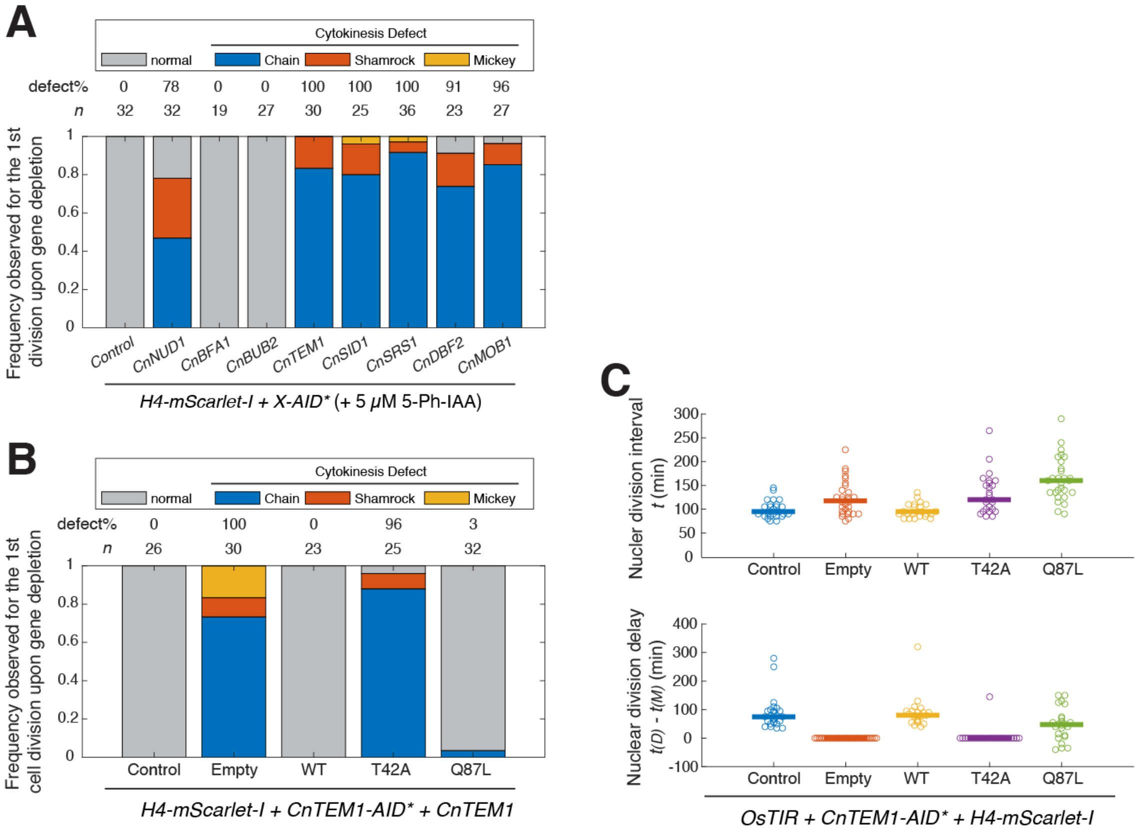


**Figure S4. The MEN/SIN pathway regulates cytokinesis in *C. neoformans***

(A) Frequency of cytokinesis defects observed for core MEN/SIN components gene depletion. Same strains and growth conditions as in Fig. 3D-E. Adding H4-mScarlet-I to the AID strains exacerbated the cytokinesis defects for *CnNUD1-AID* and *CnDBF2-AID*.

(B) Frequency of cytokinesis defects observed for CnTem1 mutants. Cn278 (Control), Cn326 (Empty, *CnTEM1-AID*), Cn407 (WT, *CnTEM1-AID* + *CnTEM1*), Cn408 (T42A, *CnTEM1-AID* + *Cntem1-T42A*), and Cn409 (Q87L, *CnTEM1-AID* + *Cntem1-Q87L*) were imaged on SC media + 2% glucose + 5 μM 5-Ph-IAA at 30 ºC and the phenotype for the first division upon 5-Ph-IAA exposure was analyzed. As predicted, the inactive GDP-locked T42A mutant displayed similar defects as empty control while the constitutively active GTP-locked Q87L mutant complemented CnTem1 depletion like WT protein.

(C) Nuclear division intervals (top) and delays (bottom) analyzed for the cells as in (B). Q87L has an increased doubling time due to a cell cycle delay like *bub2*Δ.

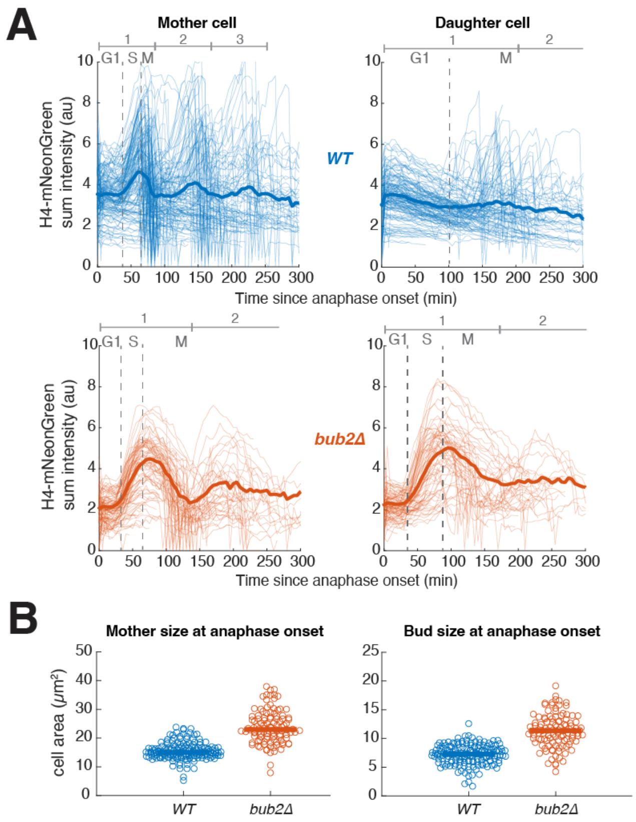


**Figure S5. *bub2*Δ cells exhibit a G2 delay.**

(A) Cell cycle timing analysis for *bub2*Δ. Cn189 (*WT*) and Cn294 (*bub2*Δ) expressing H4-mNeonGreen were grown on SC media + 2% glucose at 30 ºC and imaged every 5 min. Total H4-mNeonGreen signal were quantified for each mother and daughter cells (thin lines) following anaphase onset (estimated spindle length > 3 µm) and averaged (thick lines). The annotated G1 and S phases were estimated based on the shape of averaged H4 signal (turning point for signal increase which is assumed to reflect DNA replication). M phase was estimated based on the drop of H4 signal due to nuclear migration into the bud.

(B) Comparison of cell areas between *WT* and *bub2*Δ. Data were extracted from the same time lapse movies as (A). Cell areas were calculated using the segmented bright field images.

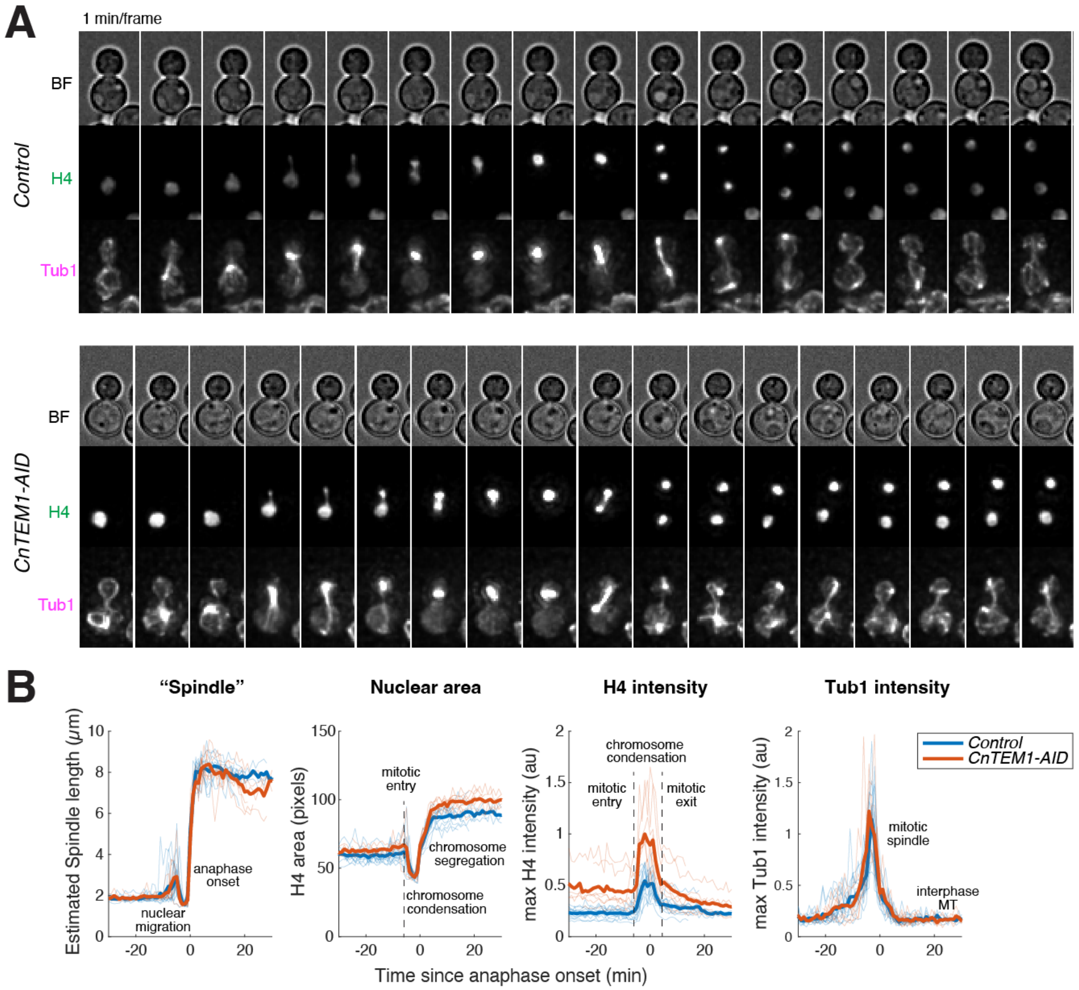


**Figure S6. The MEN/SIN pathway does not regulate mitotic exit in *C. neoformans*.**

(A) Representative time-lapse images of cells with CnTem1 depleted expressing fluorescent H4 and Tub1. Control (Cn364) and *CnTEM1-AID* (Cn350) cells expressing H4-mNeonGreen and mScarlet-I-Tub1 were grown on agarose pad made with SC media + 2% glucose + 5 μM 5-Ph-IAA at 30 ºC and imaged every minute.

(B) Comparison of mitosis timing for *CnTEM1-AID* cells. Same cells and growth conditions as (A). Spindle length (estimated by the long axis of segmented H4 signal), nuclear area (area of segmented H4 signal), H4 intensity (maximum intensity of H4 in the cell), and Tub1 intensity (maximum intensity of Tub1 in the cell) were quantified for each cell and the single cell traces (thin lines) were aligned based on the time of anaphase onset (estimated spindle length > 3 μm) and averaged (thick lines).

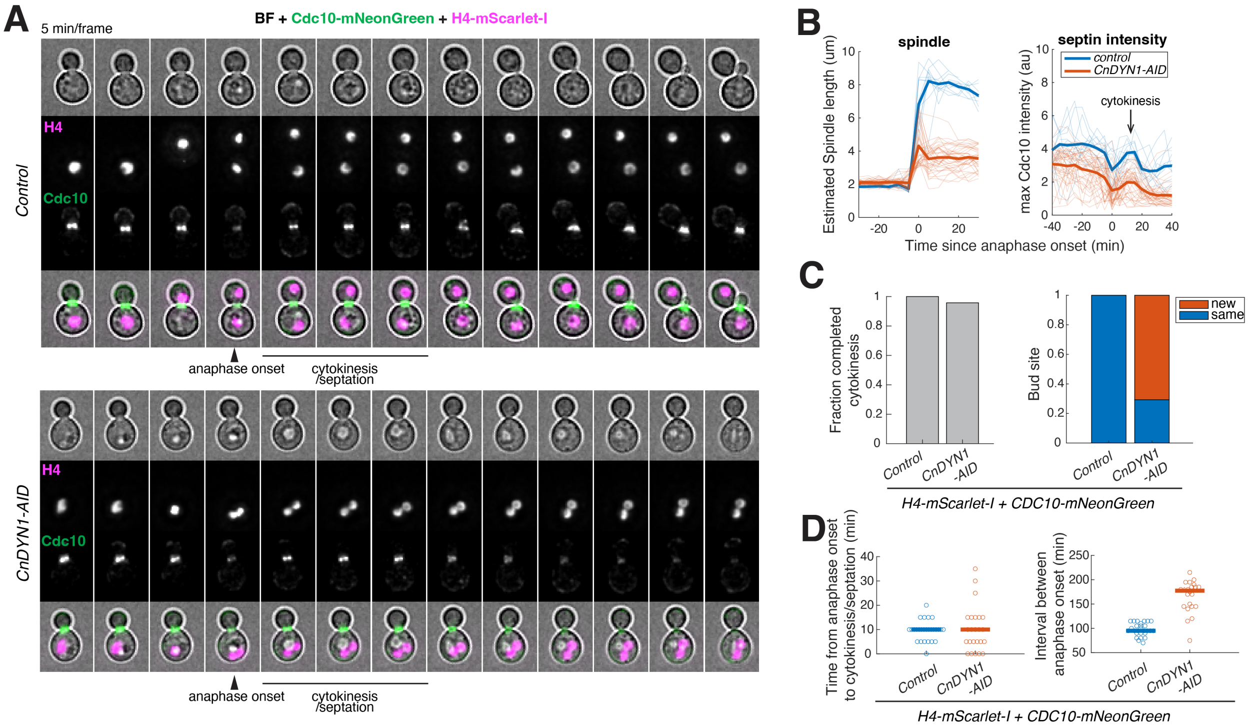


**Figure S7. Spindle position does not regulate cytokinesis in *C. neoformans*.**

(A) Representative time-lapse images of cells with CnDyn1 depleted expressing histone and septin markers. Control (Cn347) and *CnDYN1-AID* (Cn389) cells expressing H4-mScarlet-I and Cdc10-mNeonGreen were grown on agarose pad made with SC media + 2% glucose + 5 µM 5-Ph-IAA at 30 ºC and imaged every 5 min. Anaphase onset was defined as estimated spindle length (long axis of H4 signal) > 3 µm and cytokinesis/septation was labeled based on septin Cdc10 signal (peak after the drop upon anaphase onset).

(B) Comparison of mitosis and cytokinesis timing for *CnDYN1-AID* cells. Same cells and growth conditions as in (A). Spindle length (estimated by the long axis of segmented H4 signal) and septin intensity (maximum intensity of Cdc10 in the cell) were quantified for each cell and the single cell traces (thin lines) were aligned based on the time of anaphase onset (estimated spindle length > 3 µm) and averaged (thick lines). Cdc10 intensity drops upon anaphase onset which is then followed by an increase during cytokinesis (correlates with the appearance of dark line at the bud neck in bright field images).

(C) Frequency of cytokinesis (left) and bud site switching (right) for *CnDYN1-AID* cells. Same cells and growth conditions as in (A).

(D) Comparison of cytokinesis timing (left) and cell cycle timing (right) between *WT* and *CnDYN1-AID* cells. Same cells and growth conditions as in (A). Cytokinesis/septation was determined using bright field images and anaphase onset was determined using H4 signal.
